# Gene regulatory network inference with popInfer reveals dynamic regulation of hematopoietic stem cell quiescence upon diet restriction and aging

**DOI:** 10.1101/2023.04.18.537360

**Authors:** Megan K. Rommelfanger, Marthe Behrends, Yulin Chen, Jonathan Martinez, Martin Bens, Lingyun Xiong, K. Lenhard Rudolph, Adam L. MacLean

## Abstract

Inference of gene regulatory networks (GRNs) can reveal cell state transitions from single-cell genomics data. However, obstacles to temporal inference from snapshot data are difficult to overcome. Single-nuclei multiomics data offer means to bridge this gap and derive temporal information from snapshot data using joint measurements of gene expression and chromatin accessibility in the same single cells. We developed popInfer to infer networks that characterize lineage-specific dynamic cell state transitions from joint gene expression and chromatin accessibility data. Benchmarking against alternative methods for GRN inference, we showed that popInfer achieves higher accuracy in the GRNs inferred. popInfer was applied to study single-cell multiomics data characterizing hematopoietic stem cells (HSCs) and the transition from HSC to a multipotent progenitor cell state during murine hematopoiesis across age and dietary conditions. From networks predicted by popInfer, we discovered gene interactions controlling entry to/exit from HSC quiescence that are perturbed in response to diet or aging.

## Introduction

Hematopoietic stem cells (HSCs) maintain the blood system throughout life. Mammalian hematopoiesis occurs primarily in the bone marrow, where HSCs, residing in restricted niches [1, 2], give rise to multipotent progenitor cells, and subsequent lineage-restricted progenitor cell populations of various identities [3, 4]. This process is tightly controlled by a range of cell-intrinsic and extrinsic factors to avoid aberrant proliferation of stem/multipotent cells. Regulatory mechanisms controlling the lineage commitment steps during hematopoiesis are mediated primarily via gene regulatory networks (GRNs). The best-studied example of such a GRN is the pair of mutually inhibitory transcription factors (*GATA1* and *PU*.*1*) that controls commitment of hematopoietic cells into erythroid vs. myeloid (granulocyte/monocyte) lineages [5]. Another GRN motif controlling a cell fate decision is the same topology: mutual inhibition, in this case between *IRF8* and *GFI1* that control lineage commitment of the granulocyte vs. monocyte lineages [6]. Larger GRNs can also be constructed and validated, as in the case of a network detailing transcription factor interactions that regulate cell state stability [7]. However, hematopoiesis involves many other cell fate decisions — from entry into/exit from quiescence to lineage commitment — for which the core GRN interactions responsible are incomplete or unknown. Importantly, these include the early cell fate decision during hematopoiesis whereby HSCs lose their self-renewal/stemness and transition to a non-stem multipotent progenitor cell fate [8]. A combination of transcriptional and epigenetic factors are implicated in the loss-of-stemness transition from HSCs to multipotent progenitors [9, 10].

Given their required longevity [11, 12], HSCs seldom divide, and are primarily kept in a non-proliferating/quiescent state to protect against replication-induced damage [12]. Rounds of HSC division –– and thus aging –– are associated with a reduced regenerative potential and lymphopoiesis (myeloid skewing), and decreased clonal diversity [13]. These age-associated effects lead to a increase in the total number of HSCs and altered composition of the peripheral blood [12, 14, 15]. The age-related increase in HSC numbers is driven by the expansion of myeloid-biased HSCs, giving rise to disproportionately myeloid progeny [12, 16, 15]. This might partially depend on DNA damage and downstream effects [17]; more recently it emerged that epigenetic remodeling and metabolic changes also contribute to impaired HSC function [18].

Mild dietary restriction (DR), which typically consists of 60-80% of the *ad libitum* food intake by weight, has been established as a highly geroprotective intervention, leading to increases in lifespan between 20-30% in mice, which has been associated with attenuation of the aging-associated rise in inflammation, and delaying the onset of cancer and frailty [19, 20]. We have previously shown that DR in young to middle aged mice (early aging) attenuates aging-related increases in HSC numbers, and improves HSC repopulation capacity, potentially through better maintenance of stem cell quiescence [15]. The transcriptional and metabolic changes that DR induces remain largely unknown, as does the specific mechanisms by which DR alleviates aging phenotypes in HSCs. The identification of GRNs controlling cell fate decisions in hematopoiesis that are regulated through aging and/or DR could lead to an understanding of how aging drives aberrant hematopoiesis and how it may be influenced by DR.

Gene regulatory network inference seeks to determine networks of gene-gene interactions from data. Building on methods to infer GRNs from gene expression data in bulk samples (RNA-seq) [21, 22], new methods to infer gene networks have been developed in light of the higher resolution obtained by single-cell RNA-sequencing (scRNA-seq). These are built on the premise that the higher-resolution offered by decomposing bulk samples into single cells can improve GRN inference, and that the single-cell noise does not overwhelm the signal. These include methods rooted in statistical learning [23], dynamical systems theory [24], tree-based approaches [25], information theory [26, 27, 28], and time series analysis [29]. More recently, methods also consider dynamic changes to network topology itself [30]. Methods have also been introduced that make use of chromatin accessibility in addition to gene expression [31, 32, 33, 34, 35]. As a result of the variety of underlying models, these methods produce differing networks, and thus benchmarks have sought to compare their performance on both real and simulated data [36, 37, 38]. Despite their promise: it has been challenging to achieve high performance on GRN inference with single-cell data overall, especially temporally ordered data, let alone to move beyond static networks [39]. It has been shown that In some cases, the performance of pseudotime-based methods actually improves when cells are randomly ordered vs. the pseudotemporal ordering [29].

Inferences of gene regulatory interactions controlling development or stem cell differentiation in hematopoiesis and beyond have improved as the resolution of the data has dramatically increased with single-cell sequencing. More broadly, single-cell genomics approaches have shed light onto cell types [40], transition dynamics [41, 42], and cell-cell communication [43, 44] but gene expression alone may not be sufficient to accurately determine the GRNs driving cell transitions. Recent advances in experimental technologies have enabled the joint measurement of gene expression by RNA-sequencing (RNA-seq) and chromatin accessibility by assay for transposase-accessible chromatin by sequencing (ATAC-seq) in the same single cells [45, 46]. These joint multiomics data present a new opportunity to learn complex dynamic gene regulatory processes.

Here we present popInfer: network inference with pseudocells over pseudotime, a new method to infer GRNs using joint single-cell multiomic data. popInfer learns directed signed GRNs. That is, popInfer can distinguish not only the direction of interaction (regulator gene to target gene) but also the sign (activating vs. inhibitory). We tested popInfer on unperturbed hematopoiesis and on systems exposed to dietary restriction and/or aging, focusing on the dynamics of the transition from stem cells to multipotent progenitors. Through comparison with reference data gathered by chromatin immunoprecipitation assay with sequencing (ChIP-seq), we demonstrated that during homeostasis (hematopoiesis in mice fed *ad libitum*) popInfer outperforms alternative GRN inference methods that run on scRNA-seq data alone. We show that the performance of popInfer is in part derived from incorporation of pseudotime; performance drops when cells are randomly ordered.

popInfer predicted GRNs controlling the transition from HSCs to multipotent cells under different conditions. Comparative analysis of the networks revealed a core GRN governing HSC quiescence by mutual inhibition between *Mecom* and *Cdk6*. We identified a direct association between IGF signaling and the *Mecom-Cdk6* dynamics. Thus, given the increased quiescence in HSCs observed with DR at young age, we showed how HSC quiescence is controlled by IGF signaling-mediated changes in young hematopoiesis by *Mecom-Cdk6*, and how this regulation wanes as IGF signaling decreases with age.

## Results

### The stem-to-multipotent hematopoietic transition is maintained throughout lifetime and dietary perturbations

We studied early hematopoiesis in young and old mice and their response to DR via joint multiomics: concurrent sequencing of single-nucleus gene expression (snRNA-seq) and chromatin accessibility (snATAC-seq) using the 10X multiome platform (Fig. 1A). Data were generated from young (*∼* 6 months) and old (*∼* 24 months) mice. Mice were either fed *ad libitum* continually until bone marrow isolation at young (yAL) or old age (oAL), or underwent a mild dietary restriction (DR; 30% reduction in food intake by weight) for the two weeks prior to bone marrow isolation at young (yDR) or old age (oDR) to investigate early response to nutrient deprivation. Bone marrow cells were isolated and Lineage^*-*^ Sca-1^+^ cKit^+^ (LSK) cells were sorted and used for nuclei isolation and subsequent single-nucleus joint multiome sequencing (Fig. 1B). RNA-seq and ATAC-seq datasets were preprocessed (see Methods) and each modality was analyzed and integrated in our new methodology for gene regulatory network inference: popInfer.

**Figure 1:**
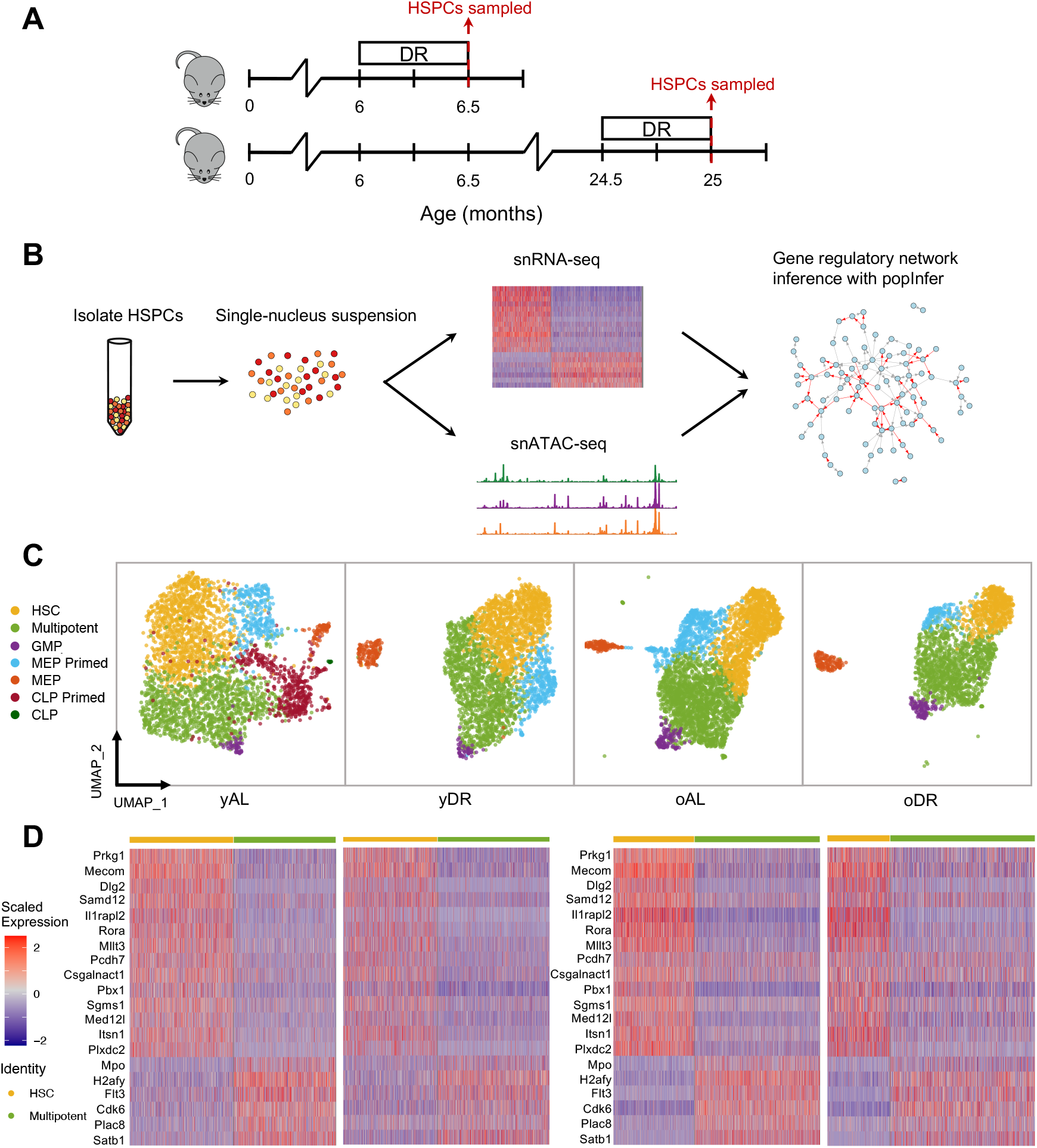
Joint multiomic data characterize hematopoiesis across diet and age. **A**. Overview of experimental design: mice undergo diet restriction (DR) at young or old age prior to sampling of hematopoietic stem and progenitor cells (HSPCs) **B**. HSPCs are sorted from the bone marrow and isolated for single nuclear (sn) RNA and ATAC sequencing, and re-integrated in a gene regulatory network inference model: popInfer. **C**. HSPCs for each experimental condition are clustered with cluster annotations made using hematopoietic marker genes. GMP: granulocyte-monocyte progenitor; MEP: megakaryocyte-erythroid progenitor; CLP: common lymphoid progenitor. **D**. Heatmaps of differentially expressed genes (top 20 ranked by log-fold change) between HSCs and multipotent progenitors; heatmaps correspond to samples above in (C). yAL: young *ad libitum*; yDR: young DR; oAL: old *ad libitum*; oDR: old DR.

Aging and diet both affect rates of cell division in the HSC pool, thereby influencing the selfrenewal and quiescence propensities of stem cells. These altered phenotypes in the HSC pool can affect hematopoiesis overall, and manifest through different frequencies of mature blood cells in the bone marrow and peripheral blood. The earliest cell fate decisions during hematopoiesis whereby HSCs may lose their self-renewal capacity during asymmetric or symmetric cell divisions when transiting to multipotent progenitor cells (MPPs) — exert influence over every cell fate decision that follows. We thus investigated how hematopoietic subpopulations change during early stages of differentiation; joint multiomic data provide a unique opportunity to identify how these transitions are transcriptionally regulated.

Unsupervised clustering via the Louvain algorithm (see Methods) identified 5-7 clusters in each condition. Via known marker genes of HSC, multipotent, and lineage-restricted early progenitor populations, cell state identities were assigned to each cluster (Fig. 1C and SI Fig 1). Across all conditions, the majority of cells belong to HSC/multipotent clusters, although we also identified lineage-primed and lineage-restricted cells for each condition (Fig. 1C).

We performed differential gene expression analysis and studied the stemness and multipotent gene expression signatures present across age and dietary conditions. We analyzed the top differentially expressed genes between HSCs and multipotent cells in yAL (ranked by log2 fold change) across samples in (Fig. 1D). Known marker genes for HSCs emerged including *Mecom* and *Mllt3*; in the multipotent cluster prior multipotent marker genes were present, including *Flt3* and *Mpo* [6, 47]. Analysis of the pattern of HSC/multipotent differential expression across all samples revealed a core hematopoietic differential gene expression pattern across all conditions. Notably, despite the relative homogeneity of the HSC/multipotent progenitor cell pool, and the clear changes that HSCs undergo throughout life, the transcriptional signature distinguishing HSCs from multipotent progenitor cells is clearly maintained throughout life and dietary restriction.

### Concurrent measurement of gene expression and chromatin accessibility enables inference of dynamic gene regulatory interactions

GRN inference methods that seek to infer gene regulatory interactions fall under two categories: time-dependent and time-agnostic methods. Several time-dependent GRN inference methods have been developed specifically for application to non-temporal (“snapshot”) data. These infer the temporal process from a population of heterogeneous single cells via trajectory inference/use of pseudotime [24, 26, 29, 48]. However, whether pseudotime values for individual genes contain useful dynamical information is highly dataset-dependent and remains an open question. Deshpande et al. [29] provided evidence that interpreting pseudotime values explicitly did not improve inference versus considering only the order of cells along the pseudotime axis. Moreover, results of GRN inference with pseudotime-ordered cells were in cases comparable to the results obtained with randomly-ordered cells.

A motivating factor in the use of pseudotime for temporal inference is the expected time lag between transcriptional changes in the regulator and those of the target (Fig. 2A). By assigning pseudotime values to cells, time lags can be modeled, even for “snapshot” data with no explicit measurement of the dynamics. Key to implementing time-series-based methods is data comprising cells that are equally spaced in time (or interpolating time-equidistant cells). That is, a reliance on the explicit use of pseudotime values in the model, which as discussed above, is an assumption that stands on shaky ground. However, inference without a temporal/pseudotemporal measure precludes the investigation of dynamic relationships between regulator genes and their targets, impeding attempts to address causality.

**Figure 2:**
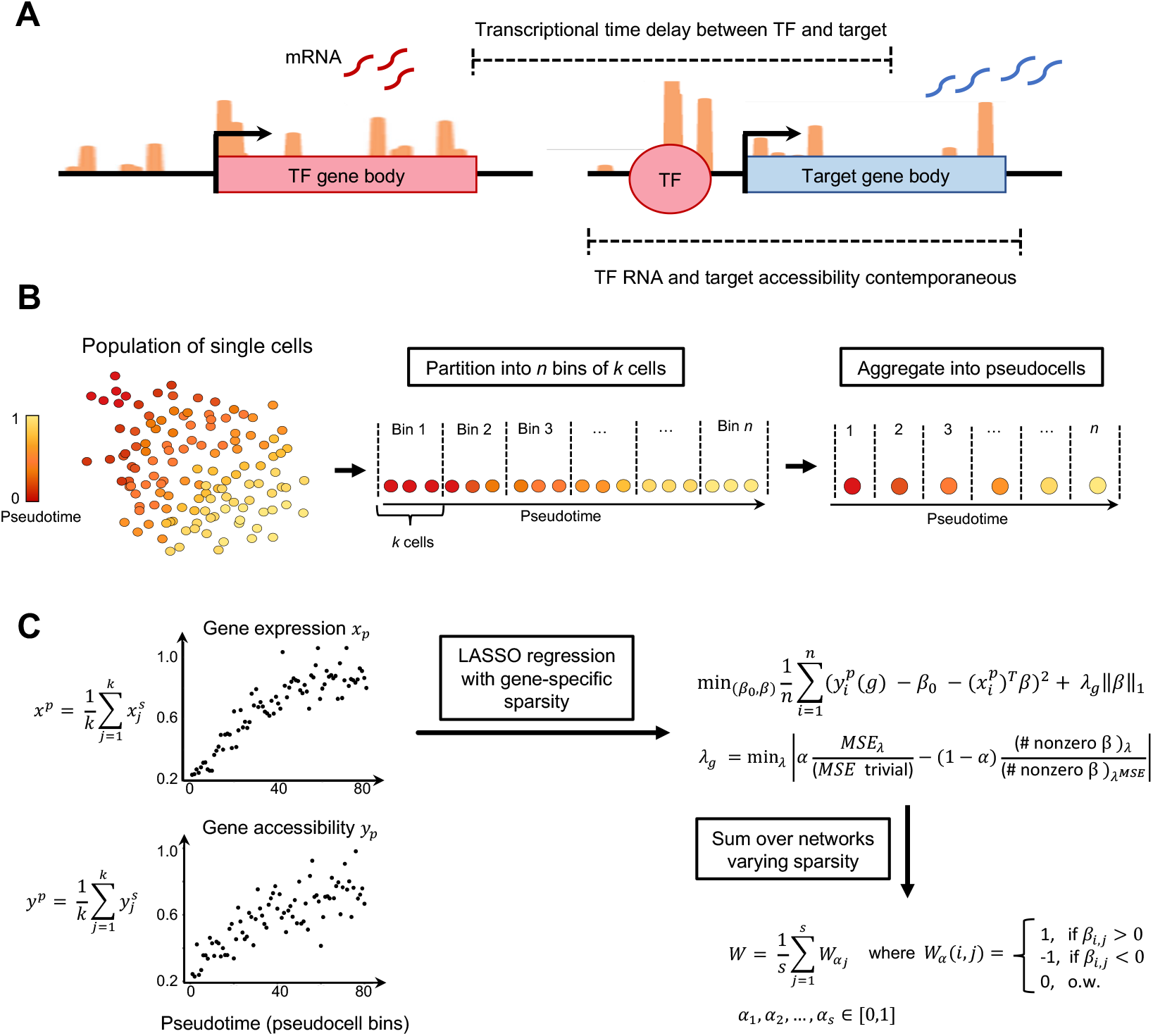
popInfer: gene regulatory network inference with multiomic data. **A**. Schematic depicting transcription factor (TF) gene expression and its regulation of a target gene. TF transcription is followed by a time delay before TF regulation can be detected in the target gene expression. TF expression is more concurrent with accessibility of the target gene. **B**. Overview of data processing for popInfer. Cells are ordered over pseudotime and binned into *n* equally sized bins of *k* cells. **B**. Overview of popInfer workflow. Cells belonging to a bin are aggregated to form a pseudocell, where 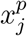 is the single-cell expression of the *j*th cell in a pseudocell bin and *x*^*p*^ is the pseudocell gene expression. 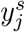 is the gene accessibility score of the *j*th cell in a pseudocell bin, and *y*^*p*^ is pseudocell gene accessibility. A LASSO regression model is run on the *n* pseudocells, where pseudocell accessibility *y*^*p*^ is predicted from pseudocell gene expression *x*^*p*^. The selection of the regularization term *λ*_*g*_ is gene-specific, governed by a tradeoff between sparsity and the mean squared error (MSE); *α* ∈ [0, 1]. *MSE*_*λ*_ is the MSE of the LASSO model for a given *λ* value and *MSE* trivial is the MSE for a trivial model with β= 0. The number of nonzero *(3* coefficients is calculated for the LASSO model for a given *λ*, and for the LASSO model that achieves the optimal MSE (*λ*^*MSE*^).

Joint multiomics offer more contemporaneous measurements of interacting genes, regulator and target, via the gene expression of the transcription factor (regulator gene) and the chromatin accessibility of the target. popInfer leverages these data to fit a regression model predicting the chromatin accessibility of the target gene from the gene expression of the regulator (Fig. 2A). While there are certainly caveats to this approach, such as the different methods used to assay each modality, these values occur more closely in time than regulator expression with target expression. Thus the assumption at the heart of popInfer is that if a TF regulates a given target, there should be a relationship between the pseudotemporal gene expression profile of the TF and the pseudotemporal chromatin accessibility of the target gene.

Single-nuclei RNA-seq and ATAC-seq data are sparse measurements in which many per-cell transcripts or peaks will be missed in sampling. This motivates our construction of pseudocells: bins of several single nuclei measurements for a given gene or ATAC-seq peak. The construction of pseudocells required means by which to partition cells into groups that were transcriptionally similar along the axis of interest (in this case stem cell differentiation). Our hypothesis is that this constitutes an appropriate opportunity to use pseudotime: as a means to order cells by their similarity along a trajectory, i.e. avoiding the need to interpret pseudotime values explicitly. Thus, we used pseudotime to order cells along a trajectory as input to partitioning into bins. Bins of cells were used to construct pseudocells, which are the input data to GRN inference by popInfer (Fig. 2B). Pseudocell gene expression (*x*_*p*_) is defined as the average expression of the cells in that bin. From the snATAC-seq data, we summarized the accessibility of a gene and its promoter region via ArchR’s GeneScoreMatrix gene accessibility scores. This score takes into account peaks that lie in the gene body and the promoter region (5kb upstream of the TSS). Pseudocell gene expression (*x*_*p*_) and gene accessibility (*y*_*p*_) scores are then defined to be the average over these values for the cells in the corresponding bin (Fig. 2C).

After constructing pseudocells with expression and gene accessibility scores, popInfer performs LASSO regression [49] using pseudocell gene expression values to predict pseudocell gene accessibility scores (Fig. 2B). For each target gene (*g*), gene-specific sparsity (*λ*_*g*_) is determined via a tradeoff between model sparsity and mean-squared error (MSE). This tradeoff is controlled by a parameter *α* ∈ [0, 1], which is fixed for all genes. When *α* = 0, *λ*_*g*_ is selected such that we obtain a trivial model whereby (β = 0. When *α* = 1, *λ*_*g*_ is selected such that we obtain a model with optimal MSE.

For a given *α*, the LASSO outputs a matrix of inferred coefficients *(3*. These values are binarized, with their sign maintained, to obtain a network matrix *W*_*α*_ (Fig. 2C). Preserving the sign of the LASSO coefficients allows popInfer to learn information regarding activation versus inhibition for interacting genes. The final output of popInfer is *W*, the averaged sum of the *W*_*α*_ for a sequence of *α* values in the range [0, 1]. Choosing *α* closer to zero produces sparser networks; larger values of *α* produce denser networks. In this study we explored the sequence of *α* values *{*0, 0.001, 0.002,…, 0.4*}*. As a result, most gene-gene pairs are assigned weights of 0. While this has the potential to inflate the false negative rate, we believe this to be overall advantageous because it is often unclear how to assign weight thresholds to select a network during GRN inference. popInfer assigned edges with zero weight are automatically be ruled out. Subsequently, a weight threshold > 0 can be set if desirable to select networks with the highest confidence edges.

### popInfer outperforms GRN inference methods based only on gene expression

To evaluate the performance of our proposed methods, we studied the results of networks inferred by popInfer and compared these with other methods for GRN inference from single-cell gene expression data. As a point of comparison, we used reference ChIP-seq data from ChIP-atlas, as defined by the benchmarking framework BEELINE [37]. For the inference methods to which we compare popInfer, we selected GENIE3 [21], PIDC [27], TENET [26], and SINCERITIES [48]. These methods have all shown strong performance under different conditions, and are built upon diverse mathematical frameworks. Benchmarking methods is inherently difficult given the challenge of identifying large sets of true positive interactions. Here we use ChIP-seq performed on hematopoietic cells from the bone marrow, which offers a set of true interactions but is an imperfect reference: since we are looking at a small subset of bone marrow cells, using a small set of variable genes, we cannot expect to recover all the true positive relationships in the reference data which measured TF-gene interactions in whole bone marrow cell populations. It is however informative to compare the relative performance of methods at recovering the highest weighted interactions from the reference dataset.

We began by studying the HSC to multipotent transition in the yAL sample. As input to GRN inference methods, we used the same set of 104 genes that were identified as differentially expressed between HSCs and multipotent progenitors (see Methods). For evaluation, we compare the results of each method via the area under the early precision-recall curve (AUEPRC). For a specified range of recall values, the AUEPRC is the area under the precision recall curve for the specified range of recall values, normalized by the total recall range. We focused on early precision-recall since we expect true networks to be sparse relative to the number of possible interactions. Under these conditions, the AUEPRC is well-suited to assess method performance (assesses whether methods perform well on the highest-weighted gene pairs). For the HSC to multipotent transition in yAL, we evaluate AUEPRC over recall values in the range [0, 0.1].

In yAL, we ran inference twice on two different sets of pseudotime values: one which was computed on expression counts that included cell cycle effects, and one which was computed on cell cycle regressed expression counts (see next section for full justification). We found that popInfer outperforms the other methods in both the cell cycle (Fig. 3A) and cell cycle regressed (Fig. 3B) iterations of inference on the yAL data. We also tested how popInfer performs when run on RNA data alone (i.e. predicting gene expression values from gene expression, as opposed to predicting gene accessibility values from gene expression; “popInfer-RNA-only”). In agreement with our logic in constructing the model (Fig. 2A), we observed a decrease in performance when using the RNA-only model, in-keeping with the assumption that the time delay between TF expression and target gene transcription incurs a cost. We found that popInfer-RNA-only performed similarly to GRN inference methods relying on scRNA-seq data alone (Figs. 3A-B).

**Figure 3:**
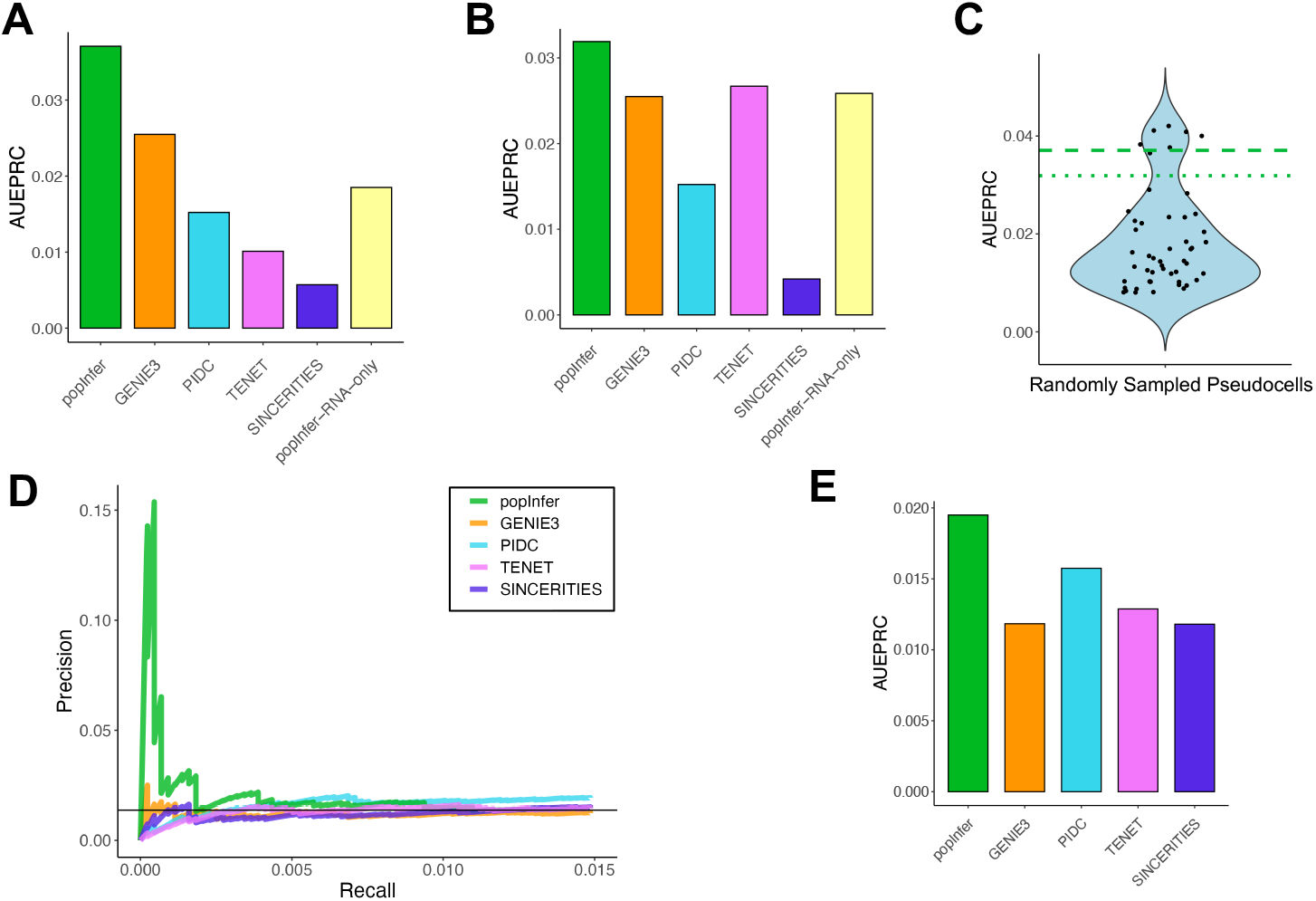
popInfer benchmark comparison with alternative GRN inference methods. **A**. Network inference evaluation on the HSC to multipotent transition in yAL for cell cycling pseudotime. Bar plots give the area under the early precision-recall curve (AUEPRC) for recall values in the range [0.0.1]. Networks were compared against a set of true positive gene interactions defined by ChIP-Atlas. **B**. Network inference evaluation with AUEPRC as for (A), with cell cycle effects regressed out. **C**. Violin plot of the area under the early precision-recall curve for yAL for popInfer, using pseudocells that were comprised of single-cells randomly sampled from cell-type clusters (as opposed to pseudotime-ordered). Green dashed line: popInfer AUEPRC from (A); green dotted line: popInfer AUEPRC from (B). **D**. Benchmarking the HSC to GMP transition in oAL; true positives from ChIP-Atlas. Early precision-recall curves for each method are shown. **E**. Bar plot of AUEPRC for the curves in (D); recall range [0, 0.015] for the HSC to GMP transition in oAL.

Next, we challenged our assumption that pseudotime was necessary for the construction of pseudocells as input to popInfer. We constructed new sets of pseudocells by randomly sampling cells from within each cluster (either HSC or multipotent) to create pseudocells. Figure 3C shows the AUEPRC of popInfer applied to these cell-type specific random pseudocells for 50 replicates. The dashed and dotted green lines are the AUEPRC value of popInfer from Fig. 3A and Fig. 3B respectively, showing that constructing psuedocells using pseudotime improves results over using cell-type resolution to create pseudocells.

Since relatively few genes comprised the set that were differentially expressed along the HSC to multipotent transition (total of 111 TF-target pairs in the reference ChIP-seq data), we also studied a hematopoietic trajectory described by a larger set of genes: differentiation from HSCs to granulocyte-monocyte progenitors (GMPs). We selected those cell belonging to HSC, intermediate multipotent, and GMP populations and studied these hematopoietic fate decisions in oAL. For features, we selected the set of differentially expressed genes between the HSC and GMP clusters in the oAL dataset, giving us 565 genes. As a result of the larger set of input genes, there were 4374 TF-target gene pairs included in the reference. We chose to use the cell cycle regressed pseudotime values for this branch of differentiation in order that cell cycle effects did not dominate over subtler gene expression/accessibility differences from the relatively few cells in the GMP cluster.

Looking at this differentiation trajectory, popInfer again outperforms other methods (Fig. 3D-E). In the early precision-recall curve, the precision values of popInfer are large relative to the other methods for small recall values, meaning that popInfer performs well at early detection of true positives (Fig. 3D). This behavior was conserved across different numbers of pseudocells and *α* sequences inputted to popInfer (SI Fig 2).

### popInfer identifies a gene regulatory network controlling HSC quiescence via IGF signaling

To study the impact of aging and dietary restriction on the HSC to multipotent cell fate transition, we ran popInfer on a set of 104 genes differentially expressed between HSCs and multipotent cells in at least one experimental condition. Gene interactions predicted by popInfer were defined as those with GRN network edge weights *>* 0.4. In the resulting networks, we considered two scenarios. Cell cycle effects can dominate pseudotime ordering, and cell cycle regression can thus be performed to avoid such cell cycle effects obscuring other regulatory interactions. However, it is not entirely possible to separate hematopoietic cell state identity with its proliferation status. Thus, we considered GRNs as predicted in each of two instances: cell cycling pseudotime, and cell-cycle regressed (non-cycling) pseudotime. For the former, no cell cycle regression is performed and cell cycle effects can be observed across cell populations (SI Fig 1B, 3); in the latter, cell cycle effects are regressed out from the snRNA-seq data before pseudotime assignment (SI Fig 1D). In both cases, diffusion pseudotime is used to construct a pseudotemporal ordering of cells (SI Fig 1C,E) [50].

Two genes stood out as hub regulators of the HSC to multipotent transition across all eight conditions (four experimental conditions; cell cycle regressed/not regressed). The first of which, Mds1 and Evi1 complex locus (*Mecom*), is a transcriptional regulator and oncogene; the second of which, cyclin-dependent kinase 6 (*Cdk6*), is a protein kinase that acts as a regulator of the cell cycle (Fig. 4A). For cell cycling pseudotime, popInfer identified *Cdk6* among the highest ranked network hubs across conditions. For cell cycle-regressed pseudotime, popInfer identified *Mecom* as a top ranked hub gene in all samples except yAL. popInfer also predicted direct interactions between *Mecom* and *Cdk6*, leading us to give closer inspection to the networks involving these genes (Fig. 4B).

**Figure 4:**
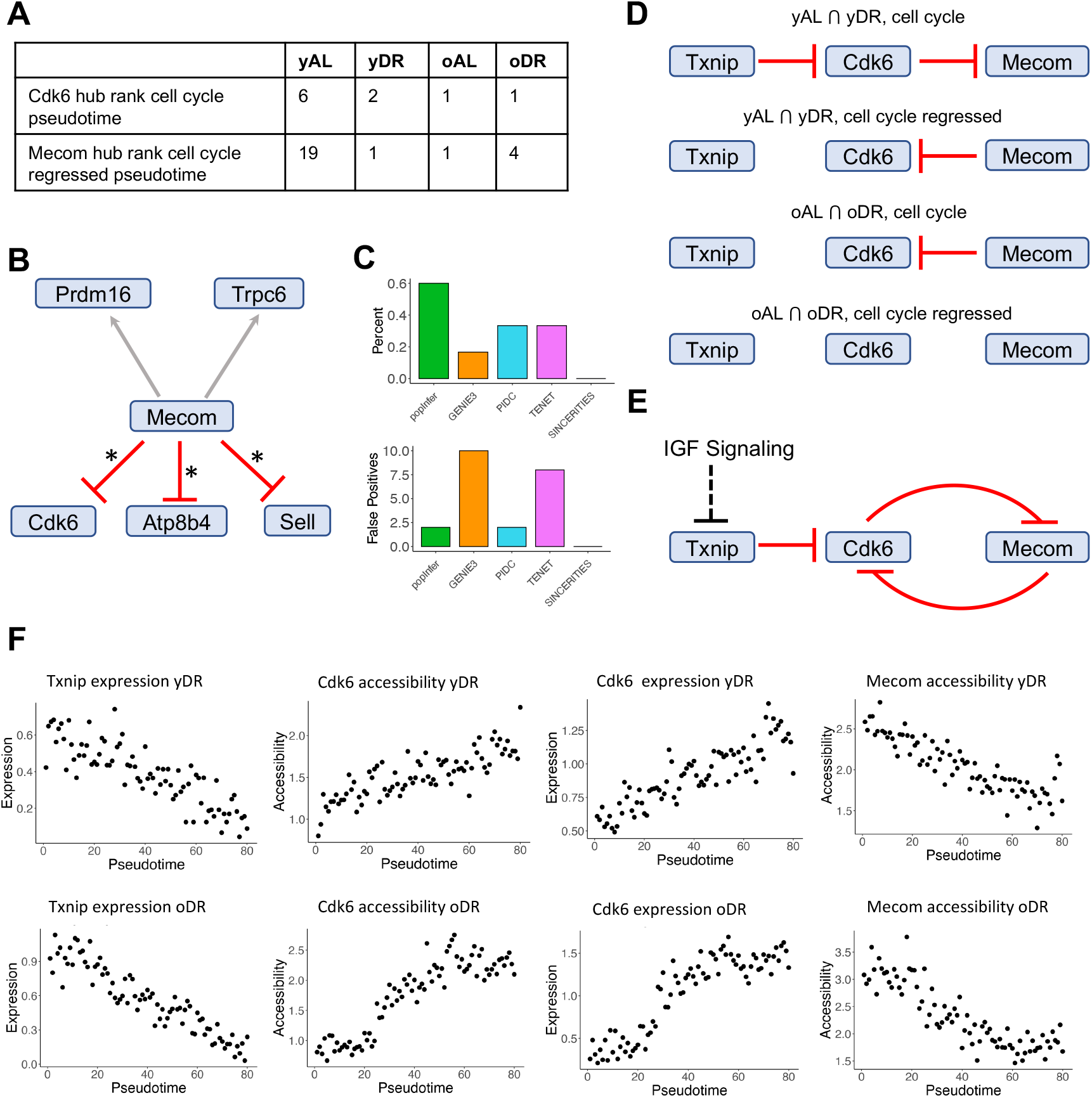
popInfer identifies regulation of HSC quiescence via IGF signaling. **A**. Table of rankings of genes *Cdk6* and *Mecom* as network hubs as inferred by popInfer. **B**. Targets of *Mecom* predicted by popInfer for yAL, using pseudotime values computed from cell cycle regressed gene expression. **C**. Bar plot of the percentage of inferred targets of *Mecom* in yAL that overlap with differentially expressed genes from [51] (top); and the number of false positives in the inferred targets of *Mecom* for each method (bottom). **D**. Consensus networks of *Cdk6* and *Mecom* between different pairs of popInfer networks. **E**. Consensus network between *Txnip, Cdk6*, and *Mecom*. Black dashed line denoting the effects of IGF1 signaling on the system is from literature. **F**. Plots of pseudocells over pseudotime (cell cycle not regressed), depicting the dynamics of *Txnip* expression, *Cdk6* accessibility, *Cdk6* expression, and *Mecom* accessibility for yDR and oDR.

As a means of comparison for popInfer predicted results, we compared them with differentially expressed genes (DEGs) that were observed upon EVI1 activation in hematopoietic progenitor cells [51]. EVI1 is a transcription factor encoded by *Evi1*, one of the alternative transcripts of *Mecom*. We found that the majority (3/5) of the target genes of *Mecom* predicted by popInfer in yAL were differentially expressed in [51], and that popInfer correctly identified the sign of the interaction in all cases (Fig. 4B). To compare popInfer results with those for other GRN inference methods, we studied how alternative methods could identify *Mecom*-associated DEGs. Using a gene interaction weight cutoff of 0.4, popInfer infers 178 network edges. For comparison, we chose the top 178 network edges for each alternative method. We found that popInfer identified EVI1-associated DEGS (as reported by Kustikova et al. [51]) with a success rate of almost twice the next best method (Fig. 4C and SI Fig 4), without sacrificing false positives (Fig. 4C lower panel).

To investigate further the relationship between the two top regulators *Cdk6* and *Mecom* during HSC to multipotent transition networks, we identified GRN interactions that overlapped between multiple experimental conditions. In young hematopoiesis, for cell cycling pseudotime, a conserved network of *Txnip* inhibition of *Cdk6*, and *Cdk6* inhibition of *Mecom* were identified (Fig. 4D). For cell cycle-regressed pseudotime, the conserved subnetwork consisted of *Mecom* inhibition of *Cdk6* (Fig. 4D). In old hematopoiesis, a conserved subnetwork consisting of *Mecom* inhibition of *Cdk6* was identified (Fig. 4D) with cell cycle pseudotime. For cell cycle-regressed pseudotime, there were no conserved interactions between control vs dietary intervention in old age. The predicted interactions are also supported by previous literature, where *Txnip* inhibition of *Cdk6* and *Mecom* inhibition of *Cdk6* have been previously reported [52, 51]. *Txnip* is a target of Insulin-like Growth Factor 1 (IGF1) of the IGF signaling pathway: an important regulator of HSC aging [53], whereby *Igf1* inhibits *Txnip*. Thus, the consensus network connects cell cycle regulation with HSC quiescence activity (popInfer predicted edges shown in red) with regulation from IGF signaling (interaction from literature shown in black) (Fig. 4E).

Analysis of the dynamics of gene expression and accessibility over the HSC to multipotent transition shed light on changes to regulatory networks with diet and upon aging. Pseudocell dynamics show that in yDR there is a strong negative correlations between *Txnip* expression and *Cdk6* accessibility, as well as between *Cdk6* expression and *Mecom* accessibility (Fig. 4F). In contrast, the dynamics of these genes in oDR are altered. While we see similar pattern of *Txnip* expression over pseudotime, its correlation with *Cdk6* accessibility is much lower due to *Cdk6* plateauing in multipotent cells both at the start and near the end of pseudotime. Concordant with its accessibility, *Cdk6* expression also plateaus at both the beginning and end of pseudotime. In contrast, *Mecom* accessibility decreases, linearly until near the end of pseudotime.

We have demonstrated how regulation of HSC quiescence is perturbed with age and diet. While *Txnip* inhibits *Cdk6* and *Cdk6* inhibits *Mecom* during hematopoiesis in young mice, these core inhibitory regulations are lost with age. Diet restriction-mediated suppression of IGF signaling increases HSC quiescence [15]. Our results described the means by which this occurs: Reduced IGF signaling upon DR leads to higher levels of *Txnip* and *Mecom*, reducing HSC cell cycle activity and driving stem cells towards a more quiescent state.

## Discussion

The ability to accurately predict GRNs that govern dynamic cell state transitions during cell differentiation and development could transform our understanding of cell fate decision-making with far-reaching consequences and applications. A persistent obstacle to achieving this goal has been the non-temporal nature of genomics data at single-cell resolution. Pseudotime has powerful applications, but is a wholly imperfect substitute for biological time, particularly in use cases for GRN inference from single-cell genomics data. Here, we proposed a solution. Joint multiomic datasets (measuring gene expression and accessibility in the same single cells) enable inference of GRNs controlling dynamic transitions via construction of pseudocells over pseudotime. We focused on the contemporaneous measurements of the gene expression of the regulatory and the chromatin accessibility of the target gene, and developed popInfer to predict directed gene regulatory interactions along with their sign (activating or inhibitory).

Through benchmarking on networks describing transitions during early hematopoiesis (HSC to multipotent progenitors, or HSC to granulocyte/monocyte progenitors), we showed that popInfer consistently outperforms alternative methods that rely on measurements of the gene expression alone. Indeed, a variant of the popInfer model that excludes chromatin accessibility data performed similarly to alternative RNA-only methods. The accuracy of networks inferred by popInfer is maintained on both small and large gene sets. Moreover, a consistent pattern emerged whereby top-weighted gene interactions predicted by popInfer were enriched with more true positive interactions (assessed by a previously published ChIP-seq reference) than alternative methods. On closer inspection of specific subnetworks, we found that for networks involving hematopoietic stem cell marker *Mecom*, popInfer predicted nearly double the fraction of true positive interactions than the next-best method. popInfer did not sacrifice sparsity to obtain these results, that is, maintaining a low false positive rate with a high true positive rate. As we have shown, jointly assaying gene accessibility and gene expression increases our power to detect gene regulatory relationships. GRN inference tools have also incorporated information regarding chromatin states in alternative ways. CellOracle constructs GRNs from single-cell gene expression and chromatin accessibility information using chromatin accessibility to construct a basis of possible GRN interactions (based on transcription factor binding and cis regulatory information [54]) and then inferring active interactions via regression using the single-cell gene expression [31]. Topic modeling methods have also been developed to characterize patterns of co-variation based on joint multiomics [55]. Argelaguet et al. [32] present a joint multiomic dataset of mouse embryonic development, and develop methods for GRN inference also making use of pseudocells [56]. Other recent approaches have employed joint multiomic data to learn GRNs from single time point data [33, 34, 35]. SCENIC+ [35] uses a large pool of putative TF-target gene interactions learnt from data to build models for GRN inference. However, despite the utility of the information gained through ATAC-seq, these GRN inference approaches still decouple the relationship between genome accessibility and target gene expression, which as we have seen (Fig. 3C), hinders our ability to learn GRNs capturing interactions during cell state transitions.

In application to early cell fate decisions during hematopoiesis, we studied hematopoietic stem and progenitor cells in young and old mice that were fed either *ad libitum* or diet restricted. For each condition, we considered two approaches: either leaving in or regressing out the effects of the cell cycle. From these analyses, we discovered a core regulatory network predicted by popInfer, governed by mutual inhibition of *Mecom* and *Cdk6*. This network regulated the transition of HSCs to multipotent cells by modulating quiescence. The network is also a target of IGF signaling, known to be reduced upon diet restriction. Thus, we have demonstrated a mechanism by which HSC quiescence is increased upon diet restriction in young age [15], concordant with decreases in *Cdk6* and increases in *Mecom*. In old age, the GRN motif regulating *Mecom* in response to IGF signaling is lost. This aligns with the finding that systemic levels of *Igf1* that influence HSC signaling decrease with aging [53]. Moreover, the expression of *Igf2bp2* — a HSC-intrinsic activator of IGF signaling — is strongly diminished in aging HSCs [57]. These HSC extrinsic and intrinsic reductions in IGF signaling fit well with the reduced IGF-dependent regulation of the *Mecom* GRN in our single cell multiome analysis of aged HSCs.

We note some limitations of popInfer. The current model does not consider genome properties (e.g. TF motifs, binding sites, or genome shape [58, 59]) in its analysis of gene accessibility, nor does it consider enhance regions or other distal regulatory elements. Incorporation of additional genome features — such as peaks from enhancer regions — into a gene accessibility score could improve specificity, although the combinatorial complexity of such a model could quickly become unwieldy. Another limitation of popInfer regards hyperparameter selection. While for the most part, popInfer-predicted networks were robust to choice of the *α* sequence, we saw that networks could be sensitive to the number of pseudocells used to bin the data. There is an inevitable trade-off: if the number of pseudocells is too small, the data become over-smoothed and transcriptional dynamic information is lost. If the number of pseudocells is too large, the data becomes too noisy and performance drops. Choice of the number of pseudocells ought to be made with care and in light of the specific dynamics under investigation. In future work use of semi-supervised methods for pseudotime could help [60]. We would also caution that if pseudocell dynamics are discontinuous or ultrasensitive (which can result from e.g. cell lineage branching in pseudotime) the assumptions underlying popInfer may no longer hold.

We have presented a model for GRN inference that uses joint multiomics in a novel way, through the integration of gene accessibility and gene expression data via regression. We anticipate wide applicability of popInfer to study cell state transitions in development and stem cell differentiation. In application to single-nucleus multiomic data describing hematopoietic stem and progenitor cells, we have elucidated a regulatory network underlying the transition from HSCs to multipotent cells. We have identified how changes in IGF signaling induced by diet restriction result in an increase in HSC quiescence via the *Mecom-Cdk6* mutual inhibitory GRN motif. Finally, we discovered that the regulation of HSC quiescence by IGF signaling is lost with aging, paving the way for future studies into mechanisms by which the loss of function in HSCs upon aging could be slowed or even reversed.

## Data and Code Availability

Single-nucleus multiome datasets are deposited and available for shared usage in the National Center for Biotechnology Information’s Gene Expression Omnibus (accession number: GSE229892). popInfer in implemented in R and available under an MIT license at: https://github.com/maclean-lab/popInfer.

## Acknowledgements

The authors would like to thank Ivonne Görlich and Marco Groth (FLI sequencing facility) for help with the multiome assay. ALM acknowledgements support from the National Institutes of Health (R35GM143019) and the National Science Foundation (DMS2045327).

## Author Contributions

MKR Conceptualization, investigation, analysis, methodology, software. M. Behrends: Investigation, analysis, methodology. YC: Investigation, analysis, methodology. JM: analysis, software. M. Bens: Investigation. LX: Methodology, software. KLR: Conceptualization, analysis, methodology, supervision. ALM: Conceptualization, analysis, methodology, software, supervision. MKR and ALM wrote the paper with input from all authors.

## Author Disclosures

The authors declare no competing interests.

## Methods

### Joint multiomics experimental methods

#### Animal experiments and housing conditions

All mouse experiments were approved by the state Government of Thuringia under the application “FLI19-009”. Male and female C57BL/6J mice were obtained from Janvier, bred in the FLI’s animal facility and kept in groups of 3-5 same sex littermates on a 12:12 hour light:dark cycle in 20-24°C and 40-60% air humidity.

The animal facilities were specific pathogen free and the mice’s cages (Tecniplast) were either individually ventilated or provided with a filter top.

#### Dietary interventions

Young (*∼* 6 months old) and aged (*∼* 24 months old) mice were randomly distributed into age- and weight-matched groups. The mice were single-housed and fed with chow prepared from commercially available powder (VRF1, SNIFF). Over the course of the first week the food intake of the individual mice was measured. At the beginning of the second week the DR group were fed once a day shortly before the onset of darkness with a food portion corresponding to 70% of the normal intake of the respective mouse. Animals were scored and weighed every other day.

#### Cell isolation and staining

The mice’s hind limbs (including hip bones joints), forelimbs and spines were dissected, cleaned, and crushed in 2% FBS using mortar and pestle. Bone marrow cells were incubated with APC-conjugated anti–c-Kit antibody, and c-Kit+ cells were enriched using anti-APC magnetic beads (MACS Milteny Biotec 130-09-855) and LS columns. C-kit positive cells were then stained with an antibody mix against mature cells (Table SI 1) for 30 minutes and overnight with a second fluorescent AB mix to stain markers used to discern different populations of HSPC (Table SI 2). The cells were sorted on an ARIA III cell sorter (BD bioscience) according to the markers in Table SI 3. 50, 000 LSK cells per mouse were sorted and cells from 2 mice per group were pooled.

#### Nuclei isolation

Cells were collected by centrifugation (10’, 300g, 4°C), the supernatant was removed and the cells were resuspended in 50*µl* 0.04% sterile filtered BSA in 1x PBS. Cells were pelleted (10’, 300g, 4°C), supernatant was discarded and nuclei isolation was conducted in accordance with the manufacturer’s protocol. In brief, 45*µl* ice-cold lysis buffer (Table SI 4) were added to the cells. Cells were pipetted up and down 3 times and incubated on ice for 3’. Subsequently 50*µl* ice-cold wash buffer (Table SI 5) were added. The samples were centrifuged for 5’ at 500g and 4°C and 95*µl* supernatant were removed. The nuclei were washed with 45*µl* ice-cold 1x nuclei buffer (Table SI 6) and centrifuged for 5’ at 500g and 4°C. 40*µl* supernatant were removed with a 100*µl* pipette, the remainder with a 10*µl* pipette. The nuclei pellet was resuspended in 7*µl* ice-cold 1x nuclei buffer.

#### 10x Multiome Protocol

1*µl* aliquot of the nuclei suspension were stained with DAPI and analysed with flow cytometry (LSRFortessa BD Bioscience) to measure final nuclei stock concentration was. Nuclei stock concentrations ranged from 3178 to 8887 nuclei/*µl*. Samples were immediately processed for scATAC-seq and scRNA-seq targeting 10,000 nuclei per sample. Samples were loaded separately onto the channels of the 10x Genomics Chromium Controller and processed with Single Cell Multiome ATAC + Gene Expression (v1 chemistry) following the standard manufacturer’s protocol (Document Number CG000338 Rev E). For ATAC-seq libraries, 7 cycles were used for the Sample Index PCR reaction and final libraries were evaluated using D5000 ScreenTape (Agilent 4200 TapeStation System) and DNA 7500 (Agilent 2100 Bioanalyzer). For scRNA-seq libraries, cDNA was amplified by 7 cycles and the total yield of cDNA was assessed on High Sensitivity DNA Assay (Agilent 2100 Bioanalyzer) resulting on average in 168 ng. A total of 13 cycles was then used for the Sample Index PCR reaction and final libraries were evaluated using D5000 ScreenTape (Agilent 4200 TapeStation System). Each type of library was pooled and sequenced using Illumina NovaSeq6000 System on SP flowcells [61]. scATAC-seq: Read 1 and Read 2 53 bp for DNA Insert, i7 8 bp for sample index and i5 24 bp for 10x Barcode and Spacer. scRNA-seq: Read 1 28 bp for 10x Barcode and UMI; Read 2 90 bp for Insert, i7 and i5 10 bp for sample index. The initial analysis with 10x Genomics Cell Ranger 2.0.0 (bcl2fastq v2.20.0.422) and corresponding pre-built mouse reference package (mm10-2020-A) estimated 2,576 to 5,833 nuclei per sample with at least 50,000 reads per nuclei.

### Joint multiomics data analysis pipeline

#### Data preprocessing and quality control

yAL, yDR, oAL, and oDR single-nucleus multiome ATAC + RNA data were processed using 10X Genomics Cell Ranger ARC (v2.0.0) mapped to the GRCm38 reference genome. Seurat (v3.2.3) and ArchR (v1.0.2) packages were used for all further analysis [62, 63].

For oAL, we began with 4754 cells. We first removed ambient RNA using SoupX [64], manually setting the contamination to 10%. Using ArchR for ATAC quality control, cells with TSS enrichment *<* 5.0 or number of fragments *<* 1200 were removed. Using Seurat for RNA quality control, we remove cells with RNA counts *>* 15000 or *<* 1500, cells with number of features *<* 1300, or cells with mitochondrial percentage *>* 20%. After this initial quality control, 4082 cells remained. Lastly, we ran Doublet Finder [65] with 3% doublet formation rate and PC neighborhood size *pK* = 0.29, removing 122 doublets and leaving us with a final set of 3960 cells (Fig. 1A).

For oDR, we began with 2576 cells. We first removed ambient RNA using SoupX, manually setting the contamination to 10%. Using ArchR for ATAC quality control, cells with TSS enrichment *<* 8.0 or number of fragments *<* 2000 were removed. Using Seurat for RNA quality control, we remove cells with RNA counts *>* 10000 or *<* 1000, cells with number of features *<* 500, or cells with mitochondrial percentage *>* 35%. After this initial quality control, 2299 cells remained. Lastly, we ran Doublet Finder with 3% doublet formation rate and PC neighborhood size *pK* = 0.17, removing 69 doublets and leaving us with a final set of 2230 cells.

For yAL, we began with 4185 cells. We first removed ambient RNA using SoupX, manually setting the contamination to 10%. Using ArchR for ATAC quality control, cells with TSS enrichment *<* 5.0 or number of fragments *<* 2000 were removed. Using Seurat for RNA quality control, we remove cells with RNA counts *>* 9000 or *<* 1000, cells with number of features *<* 700, or cells with mitochondrial percentage *>* 25%. After this initial quality control, 3423 cells remained. Lastly, we ran Doublet Finder with 3% doublet formation rate and PC neighborhood size *pK* = 0.29, removing 103 doublets and leaving us with a final set of 3320 cells.

For yDR, we began with 3616 cells. We first removed ambient RNA using SoupX, manually setting the contamination to 10%. Using ArchR for ATAC quality control, cells with TSS enrichment *<* 5.0 or number of fragments *<* 3000 were removed. Using Seurat for RNA quality control, we remove cells with RNA counts *>* 9000 or *<* 700, cells with number of features *<* 500, or cells with mitochondrial percentage *>* 40%. After this initial quality control, 3146 cells remained. Lastly, we ran Doublet Finder with 3% doublet formation rate and PC neighborhood size *pK* = 0.27, removing 94 doublets and leaving us with a final set of 3052 cells.

#### Hematopoietic cell subpopulation clustering and peak calling

For all datasets, RNA counts were transformed by regressing out *G*2*M* and *S* phase cell cycle scores (from the CellCycleScoring Seurat function) using the Seurat function SCTransform prior to clustering (Fig. 1B,D). In all datasets, the first 30 principal components were used for louvain clustering and Uniform Manifold Approximation and Projection (UMAP) in Seurat. Clustering resolution was set to 0.25 for *oAL* and *oDR*, 0.35 for *yAL*, and 0.31 for *yDR*, resulting in 6 distinct clusters in oAL, 7 distinct clusters in oDR and yAL, and 5 clusters in yDR. Corresponding cell types were annotated using canonical hematopoietic cell type markers. For peak calling, all four datasets were used to create a single ArchR object. Using the clusters defined using the RNA, the two smallest clusters in both *oAL* and *oDR* were removed prior to peak calling and the smallest cluster in *yDR* was removed prior to peak calling. Pseudo-bulk replicates were made in ArchR using the addGroupCoverages function with maxReplicates set to 10 and groupBy set to the samples and clusters defined using the RNA data. The pseudo-bulk replicates were used for peak calling via the addReproduciblePeakSet ArchR function.

### popInfer model for gene regulatory network inference

#### Feature selection for network inference

To select features to use as input to the gene regulatory network inference model, we first identified genes that were differentially expressed between the HSC and multipotent progenitor clusters for each of the four samples using Seurat’s FindMarkers function. We defined a gene to be differentially expressed if it’s adjusted *p*-value was less than 0.05. We then took the union over these four sets of DEGs, giving us 107 genes. From this set of genes, we removed three genes that weren’t found in the reference chromatin annotation. The remaining 104 genes were used as the input features to the network inference method. For the transition from HSCs to GMPs in oAL, we used the 565 differentially expressed genes between the HSC and GMP clusters as the input features.

#### Pseudotime ordering and pseudocell construction

On each dataset, DPT [50] was applied to the RNA assay to assign pseudotime values. The root cell of DPT was set to be the cell with highest expression of the sum of Mecom and Mpl, two canonical HSC markers. For yAL, the root cell was defined as the cell with the fourth highest expression of the sum of Mecom and Mpl because the top three ranking cells either did not lie in the HSC cluster or were on the boarder between the HSC cluster and another cluster in the UMAP. The expression of the top 2000 variable features were used as input to DPT.

For each dataset, two different pseudotime assignments were produced: one which was computed on expression counts that included cell cycle effects, and one which was computed on cell cycle regressed expression counts (Fig. 1C,E). DPT [50] was therefore applied twice to each sample. The first iteration of DPT was run on counts that were transformed after quality control using the Seurat function SCTransform. The second iteration of DPT was run on counts were transformed by regressing out *G*2*M* and *S* phase cell cycle scores (from the CellCycleScoring Seurat function) using the Seurat function SCTransform.

For each dataset and for each pseudotime assignment, we constructed pseudocells. To do so, for each dataset, cells were first ordered along pseudotime. Cells are then partitioned into similarly sized pseudocell bins where the first *n* mod 𝓁 bins contain 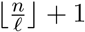 cells and the remaining *n -* (*n* mod 𝓁) bins contain 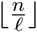 cells. Pseudocell expression (*x*^*p*^) was defined as the average of the expression of the *k* cells within it’s corresponding bin:

average of the expression of the k cells within it’s corresponding bin:

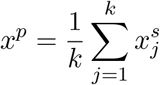

where 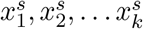 are the expression values of the cells in the pseudocell bin.

For constructing pseudocell gene accessiblity scores, we used ArchR’s GeneScoreMatrix function. Using the default parameters, gene scores are computed using genes in the gene body and 5kb upstreamm of the TSS. Pseudocell accessibility (*y*^*p*^) was defined as the average of the expression of the *k* cells within it’s corresponding bin:

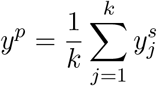

where 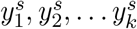 are the ArchR GeneScoreMatrix accessibility scores of the cells in the pseudocell bin for pseudocell *p*.

### Pseudotime-ordered pseudocell (POP) inference model

From our pre-processing, we have *n* pseudocells, for which we have pseudocell expression scores 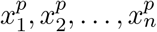 and pseudocell gene accessibility scores 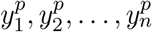. From our feature selection, we have genes *G*. For each gene *g*∈ *G*, we run a lagged LASSO regularized linear regression model using glmnet [66]:

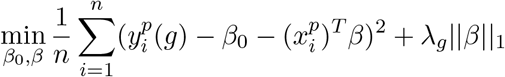

Where λ_*g*_ is selected via the optimization:

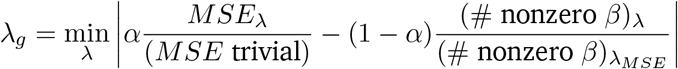

where α ∈ [0, 1], *MSE*_*λ*_ is the MSE of the LASSO model for .*>, MSE* trivial is the MSE when we have a trivial model ((β= 0), (# nonzero (β) λ is the number of nonzero coefficients of the LASSO model for a given. λ value, and. λ^*MSE*^ is the value of λ for which the LASSO model achieves optimal MSE. We implement gene-specific sparsity parameters (λ_*g*_) because we found that using the same *λ* value across all genes resulted in inconsistent levels of sparsity (relative to the sparsity in the optimal MSE model for that gene). The optimization we defined thus allows for a better control over global sparsity of the model. We evaluate this optimization for a given *α* value by running glmnet for 200 λ values and solving [66].

After running for a fixed *α* value, we define a *G* × *G* output matrix,

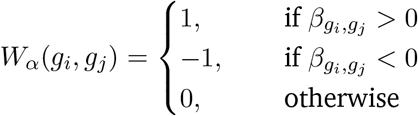

for all *g*_*i*_, *g*_*j*_ ∈ *G, i* ≠ *j*. Finally, we run for a sequence of *α* values *α*_1_, *α*_2_,…, *α*_*s*_ ∈ [0, 1] and define the final output of popInfer to be the weight matrix:

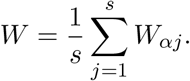

In this work, we use the sequence of *α* values: *α 2 {*0.001, 0.002,…, 0.4*}* with the exception of the oAL GMP branch testing, for which we used the sequence of *α* values: *α* ∈*{*0.001, 0.002,…, 0.6*}*. We select *α* values closer to 0 because we expect GRNs to be sparse and according to our defined optimization problem, values of *α* closer to zero will be more sparse. Averaging over the results of a sequence of *α* values as opposed to only running for a single value of *α* is helpful in a number of situations, such as when a gene has a nonzero value of βonly a few times by chance or if a gene interaction has inconsistent predicted sign (activating/inhibiting) for different *α* values. In both of these examples, sporadic or inconsistent relationships between a predictor and target gene will be mitigated by running for a sequence of *α* values.

## Notes

https://github.com/maclean-lab/popInfer

### Competing Interest Statement

The authors have declared no competing interest.

