## Supplementary Figures for "Gene regulatory network inference with popInfer reveals dynamic regulation of hematopoietic stem cell quiescence upon diet restriction and aging"

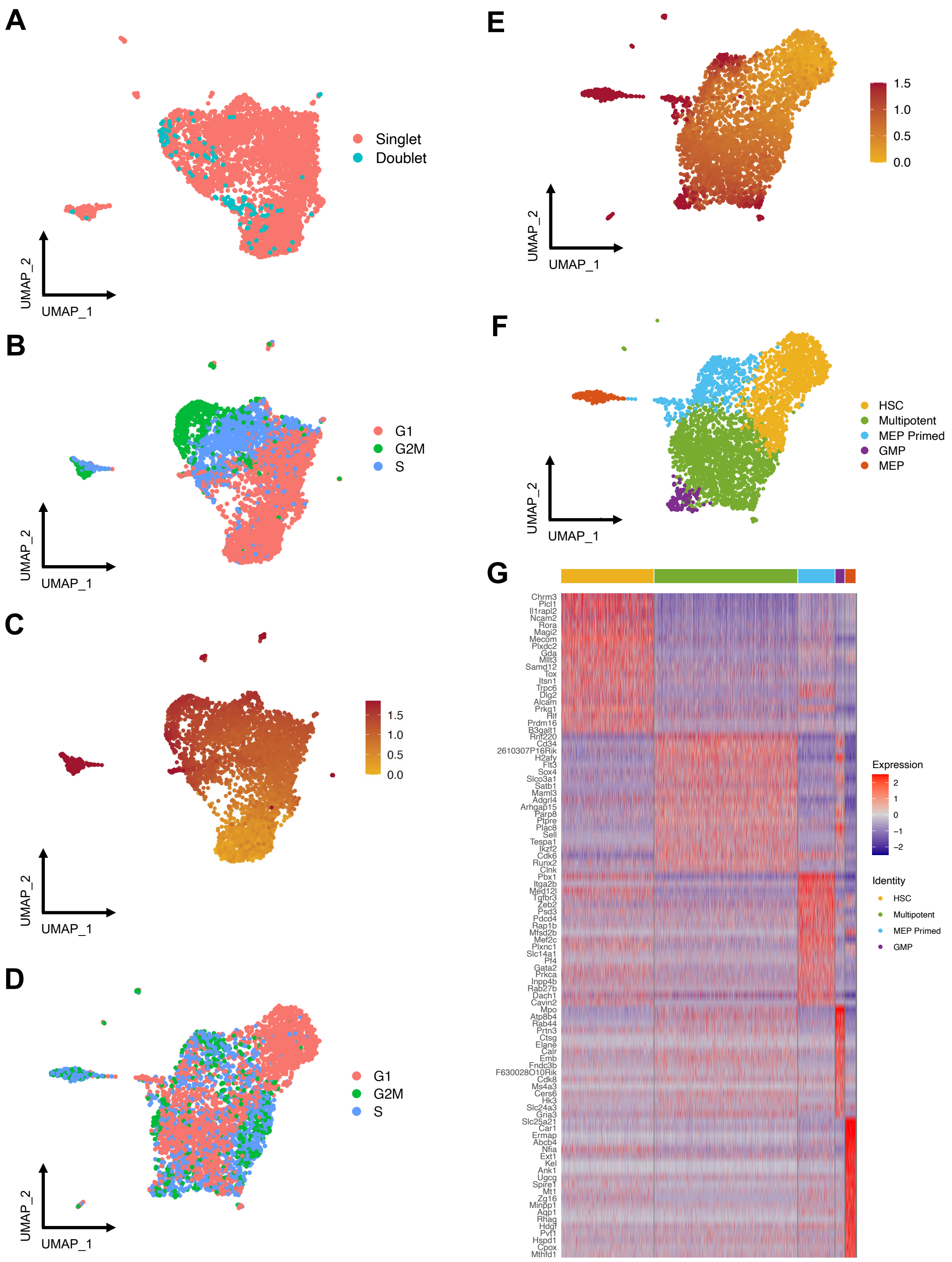

Figure SI 1: Overview of single-nucleus multiome data preprocessing workflow in oAL. (A) UMAP of DoubletFinder identified singlets and doublets in oAL. (B) After removal of doublets, UMAP of oAL data where cells are colored by cell cycle phase. Cell cycle effects appear significant, leading to cells clustering by phase. (C) UMAP of oAL data colored by pseudotime value. Pseudotime values greater than 2.0 were set to be 2.0 in order to clearly see the color gradient on the plot. (D) UMAP of oAL data after cell cycle effects are regressed. (E) UMAP of cell cycle regressed oAL data colored by pseudotime value. Pseudotime values greater than 1.5 were set to be 1.5 in order to clearly see the color gradient on the plot. (F) Clustering and cell type annotation of the cell cycle regressed oAL UMAP with clustering resolution set to $0.25$. (G) Heatmap of top differentially expressed genes by cluster.

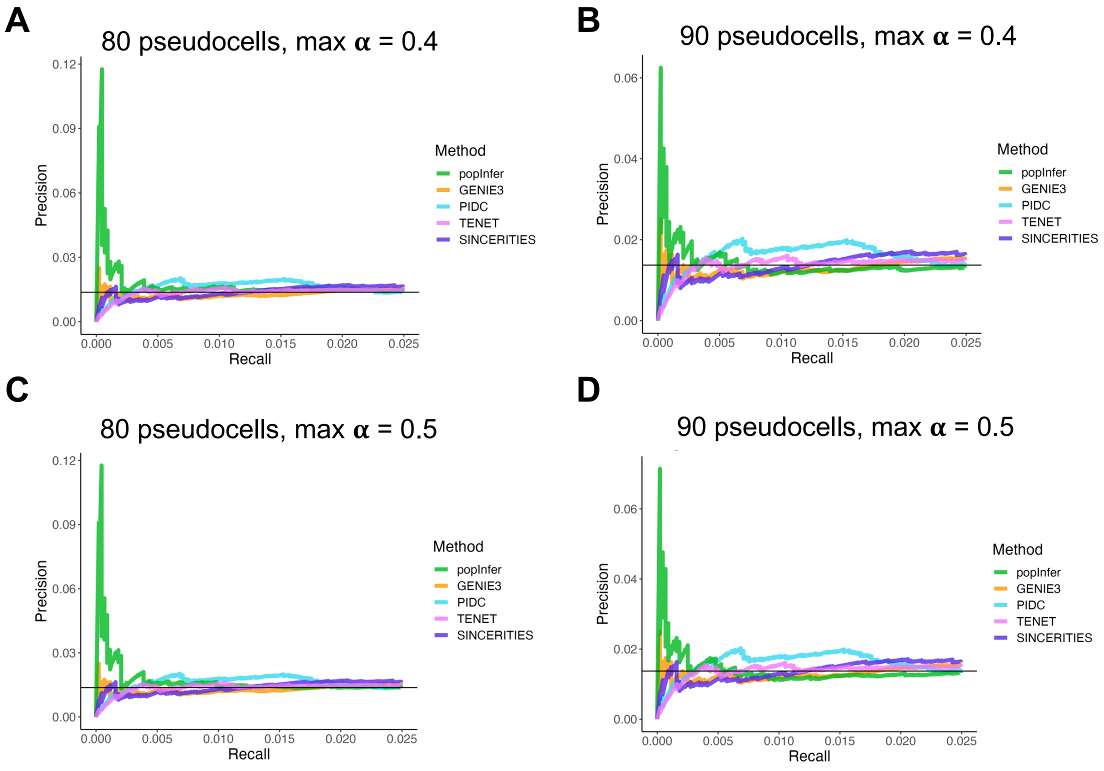


Figure SI 2: Early precision-recall curves for the HSC to GMP transition in oAL for different numbers of pseudocells and $\alpha$ sequences: (A) 80 pseudocells, $\alpha \in \{0, 0.001, 0.002, \ldots, 0.4\}$, (B) 90 pseudocells, $\alpha \in \{0, 0.001, 0.002, \ldots, 0.4\}$, (C) 80 pseudocells, $\alpha \in \{0, 0.001, 0.002, \ldots, 0.5\}$, (D) 90 pseudocells, $\alpha \in \{0, 0.001, 0.002, \ldots, 0.5\}$.

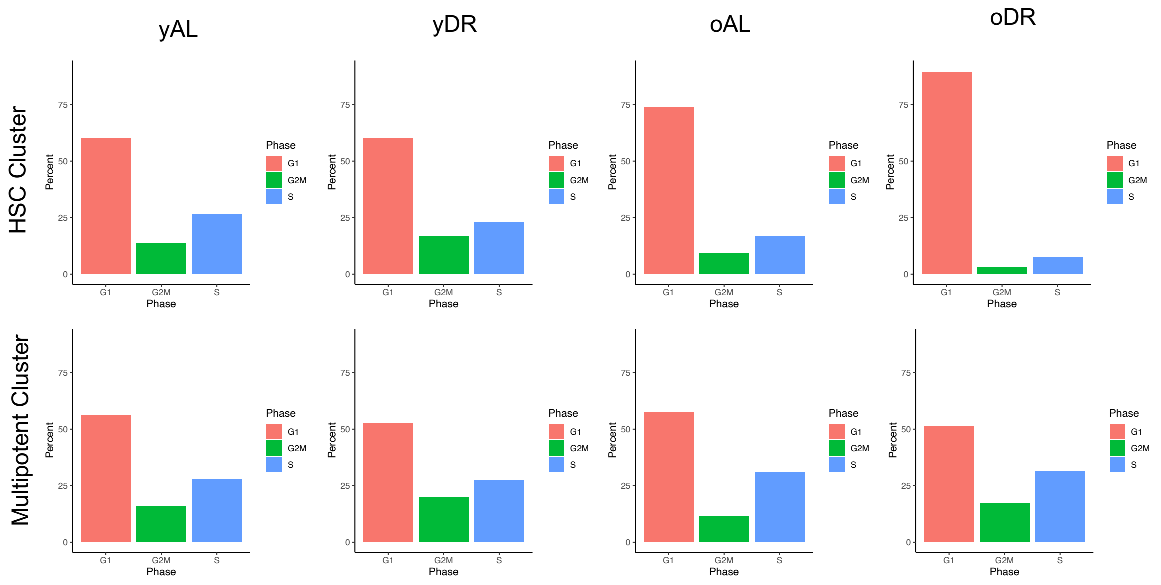


Figure SI 3: Percent of HSC and multipotent cells in each cell cycle phase by sample.

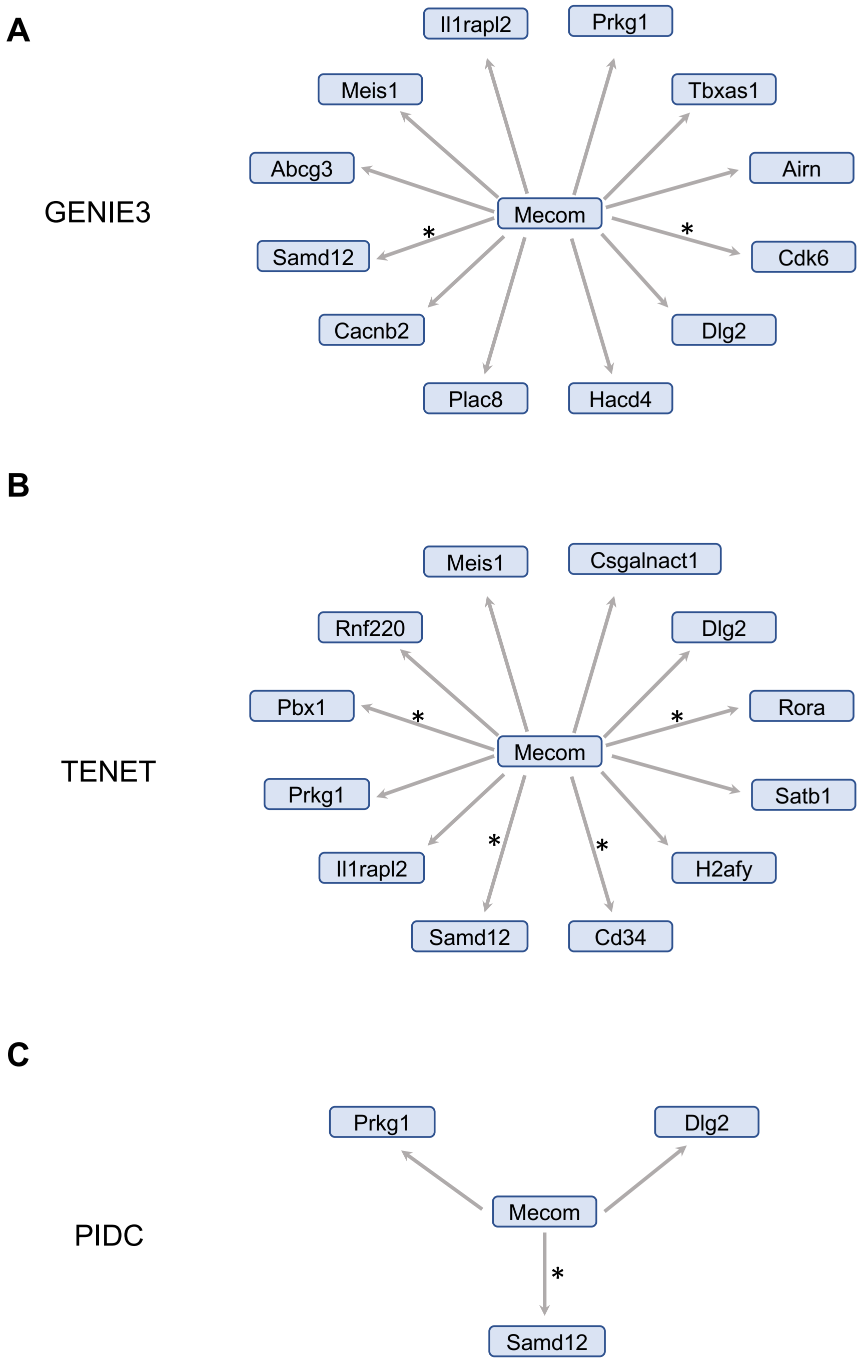


Figure SI 4: Subnetworks of yAL *Mecom* targets inferred by (A) GENIE3, (B) TENET, and (C) PIDC. Edges marked with black stars correspond to target genes that were differentially expressed upon *Evi1*( encoded by *Mecom*) activation.
