## Supplementary Tables for "Gene regulatory network inference with popInfer reveals dynamic regulation of hematopoietic stem cell quiescence upon diet restriction and aging"

### Fluorescent antibodies used for cell staining

| Reagent | Company | Catalogue Number | Clone Number |
|---|---|---|---|
| Biotin TER 119 | Bio Legend | 116204 | TER-119 |
| Biotin GR1 | Bio Legend | 108404 | RB6-8C5 |
| Biotin CD11b | Bio Legend | 101204 | M1/70 |
| Biotin B220 | Bio Legend | 103204 | RA3-6B2 |
| Biotin CD4 | Bio Legend | 100508 | RM4-5 |
| Biotin CD8a | Bio Legend | 100704 | 53-6.7 |

Table SI 1: Mature cell fluorescence AB mix to stain lineage-markers.

| Reagent | Company | Catalogue Number | Clone Number |
|---|---|---|---|
| CD150 Brilliantviolet 605 | Bio Legend | 115927 | TC15-12F12.2 |
| c-kit APC | Bio Legend | 105812 | 2B8 |
| SCA1 (Ly-6A/E) PE | Bio Legend | 108108 | D7 |
| Streptavidin APC-Cy7 | Bio Legend | 405208 | - |
| CD34 FITC | Invitrogen | 11-0341-85 | RAM34 |
| CD16/32 PE-Cy7 | Bio Legend | 101318 | 93 |

Table SI 2: HSPC fluorescence AB mix.

| Cell type | Marker/ definition |
|---|---|
| LSK | Lin-/c-kit+/Sca1+ |
| MEP | Lin-/c-kit+/Sca1-/FcR-/CD34- |
| CMP | Lin-/c-kit+/Sca1-/FcR-/CD34+ |
| GMP | Lin-/c-kit+/Sca1-/FcR+/CD34+ |
| MPP | Lin-/c-kit+/Sca1+/CD34+ |
| HSC | Lin-/c-kit+/Sca1+/CD34-/ CD150+ |

Table SI 3: Markers of HSPC populations

| | Stock | Final | Volume for 1ml in $\mu$l |
|---|---|---|---|
| 20x Nuclei Buffer | 20x | 1x | 50 |
| DTT | 1000mM | 1mM | 1 |
| RNase inhibitor | 40U/$\mu$l | 1U/$\mu$l | 25 |
| Nuclease free water | - | - | 924 |

Table SI 4: Diluted nuclei buffer

| | Stock | Final | Volume for 4ml in $\mu$l |
|---|---|---|---|
| Tris-HCl pH 7.4 | 1M | 10mM | 40 |
| NaCl | 5M | 10mM | 8 |
| MgCl2 | 1M | 3mM | 12 |
| BSA | 10% | 1% | 400 |
| Tween-20 | 10% | 0.1% | 40 |
| DTT | 1000mM | 1mM | 4 |
| RNase inhibitor | 40U/$\mu$l | 1U/$\mu$l | 100 |
| Nuclease free water | - | - | 3400 |

Table SI 5: Wash buffer

| | Stock | Final | Volume for 2ml in $\mu$l |
|---|---|---|---|
| Tris-HCl pH 7.4 | 1M | 10mM | 20 |
| NaCl | 5M | 10mM | 4 |
| MgCl2 | 1M | 3mM | 6 |
| Tween-20 | 10% | 0.1% | 20 |

Table SI 6: Lysis buffer
